# A field-deployable eDNA metabarcoding workflow including *de novo* reference assembly for characterizing understudied biodiversity hotspots

**DOI:** 10.1101/2025.09.14.676136

**Authors:** Jesse Erens, Christopher Heine, Stefan Lötters, Henrik Krehenwinkel, Andrew J. Crawford, Luis Alberto Rueda-Solano, Amadeus Plewnia

## Abstract

Field-deployable DNA metabarcoding offers a transformative approach to biodiversity research and monitoring, yet its application remains limited due to technical constraints and a lack of reference data in poorly studied ecosystems. Combining isothermal Recombinase Polymerase Amplification (RPA) and Oxford Nanopore sequencing, we introduce a two-step approach that uses non-invasive species barcoding to directly generate reference sequences for use in environmental DNA (eDNA) metabarcoding, and enable real-time, PCR-free, and cost-effective molecular assessment of ecological communities in the field. Using an endemic and understudied tropical amphibian assemblage as a model, we demonstrate the practicality and versatility of this novel workflow. *De novo* generation of a reference sequence library significantly improved the accuracy and taxonomic resolution of sequence assignments from eDNA samples, particularly on the species level, in turn allowing a characterization of fine-scale patterns in community composition. Beyond generating new RPA-compatible amphibian metabarcoding primers, our results show that combining field-based eDNA metabarcoding with the offline assembly of a local reference database can bridge data gaps in molecular biodiversity monitoring, providing a scalable solution for real-time biodiversity assessments in data-deficient ecosystems. This workflow paves the way for broader deployment of molecular tools in global biodiversity hotspots – particularly in remote and resource-limited tropical regions – to directly contribute critical baseline data, and support conservation efforts in regions where they are most urgently needed.

## Introduction

In the face of rapid and ongoing biodiversity decline, improved access to field-deployable, cost-effective, and non-invasive molecular species detection tools is of primary importance (Kühl et al., 2020; Theissinger et al., 2023; Plewnia et al. 2025a). Since global resources for biodiversity conservation remain limited, it becomes essential to be able to prioritize species and communities to achieve timely and effective conservation action (Wilson et al. 2006). Among the key limitations that currently hamper our efforts to preserve wildlife populations, however, are a lack of knowledge on how species are exactly distributed and how many taxa occur within a given area (Bini et al. 2006; Brito 2010). These shortfalls are most severe in tropical regions, the very areas that harbour the highest levels of biodiversity but often lack sufficient resources for comprehensive research and conservation (Collen et al., 2008). Hence, there is an acute need to make molecular biodiversity monitoring methods more accessible and cost-efficient to enable their wider application in understudied and hyperdiverse tropical ecosystems.

Community metabarcoding is a rapidly advancing toolkit employed in the standardized assessment of biodiversity. Its application enables comprehensive assessments of biodiversity change and community composition across scales ranging from microbial communities to entire biomes (Compson et al., 2020; Creer et al., 2016; Gillespie et al. 2023). The development of portable and lower-cost laboratory equipment and high-throughput sequencing technologies is currently democratizing molecular biodiversity monitoring (Hatfield et al., 2023; Krehenwinkel et al., 2019; LaBarre et al., 2011; Ríos et al., 2012; Plewnia et al. 2025b), making the metabarcoding toolkit increasingly accessible to conservationists, practitioners and researchers in remote, resource-limited settings. Presently, rapid DNA extraction can be combined with Recombinase Polymerase Amplification (RPA) – an isothermal method not requiring a thermal cycling machine – to build libraries from environmental samples that can be sequenced on site with a portable Oxford Nanopore Technology sequencing device (hereafter: ‘Nanopore’) (Plewnia et al. 2025b). However, field-deployable metabarcoding is still in its infancy, and several significant limitations hinder its broader application for eDNA studies.

First, constraints remain due to the specific requirements for RPA primer design (Li et al., 2019; Lobato & O’Sullivan, 2018). RPA performs best with long primers (∼30 bases), which are prone to artefact formation (Lobato & O’Sullivan, 2018), thus challenging the design of such long primers for higher taxa that commonly share only short conserved regions. For environmental DNA metabarcoding, primer design is further limited to the few, mostly mitochondrial markers, for which broad sequence libraries are available to serve as a reference for taxonomic assignment. Nevertheless, RPA has shown greater tolerance to sequence mismatches at primer binding sites (Daher et al., 2015), increasing its potential applicability across closely-related species. Thus, the emergence of RPA introduces both new challenges and opportunities, and an experimental approach is still required to assess the compatibility of existing PCR primers. As such, general guidelines and analytical tools for adapting PCR primers for RPA-based metabarcoding are still largely lacking.

Second, the incompleteness and inaccuracy of DNA reference databases, particularly for the tropical sampling gap, presently limits molecular taxonomic identification of Operational Taxonomic Units (OTUs) from environmental samples (Hughes et al., 2021; Jetz et al., 2012). The development of portable sequencing devices, with the Nanopore MinION being the most widely used platform, now allows high-throughput DNA barcoding of hundreds of specimens with minimal equipment in the field (Krehenwinkel et al., 2019; Pomerantz et al., 2022). Complementing field-deployable eDNA metabarcoding with simultaneous reference database creation from targeted species-specific barcoding could substantially increase taxonomic accuracy, thus accelerating the real-time characterization of ecological communities in the field.

In this paper, we address these limitations and expand the RPA-based metabarcoding workflow. We do this by (1) nesting a novel environmental DNA (eDNA) metabarcoding primer set within a long-read specimen barcoding marker. This enables fully isothermal, field-deployable community metabarcoding from environmental samples with species identification being guided by (2) a reference database that is *de novo* assembled in the field using non-invasive DNA sampling methods and the same sequencing workflow as used for environmental samples. We demonstrate the real-world application of this novel approach through the molecular characterization of an understudied tropical montane amphibian community of high conservation relevance. By further expanding open-source protocols for RPA primer development for metabarcoding and local, offline reference database assemblies, we facilitate the application of field-based metabarcoding to advance global efforts in biodiversity assessment and conservation.

## Methods

### RPA primer design

Our eDNA assay targets the 16S ribosomal RNA gene, one of the most widely used genetic markers for barcoding amphibians (Kocher et al., 1989; Palumbi et al., 2002; Salducci et al., 2005; Vences et al., 2005), making it highly suitable for eDNA metabarcoding (Sakata et al., 2021). We started with various previously utilized PCR primers designed for different amplicons within 16S and further modified primers by manually adding bases to one or both ends to increase their potential for RPA-compatibility. As the foundation for primer design, we used the recent 16S alignment of Portik et al. (2023), which includes sequence data for 4,890 anuran species. As we visually identified misaligned regions in the dataset, we re-aligned all 16S sequences using MAFFT (Katoh and Standley, 2002) and removed insertions and divergent regions with CIAlign (Tumescheit et al. 2022) as well as regions with >50 % gaps in all sequences using trimAL (Capella-Gutiérrez et al. 2009). We then aligned existing primers (Palumbi et al. 2002; Sakata et al. 2021) and visually screened for additional conserved regions and conserved positions bordering the existing primer binding sites. Based on the identified conserved positions, we then elongated existing primers to a total of ∼25-30 bases while minimizing the number of degenerate bases. In addition to the ‘universal’ 16S primer 16Sbr (Palumbi et al. 2002), we designed a total of 24 new candidate RPA primers spread across six different binding sites along the 16S gene for further testing with RPA (Table S1), with various target amplicon lengths (Figure S1). We assessed the RPA functionality of 77 primer combinations using amphibian DNA isolates from tissue samples (given material constraints, we could not test all primer combinations). This yielded 29 pairs that showed visually consistent amplification based on gel electrophoresis, which were selected for further testing amplification success from eDNA samples (Figure S2). The five candidate primer pairs with the strongest visual amplification of eDNA samples based on gel electrophoresis were taken for further Nanopore indexing and a test MinION sequencing run (Figure S2). For dual barcoding of each sample, we added 20-base indexes adopted from Gajski et al. (2024) to the 5’ end of each selected primer. Selected primers (including indexes) were checked *in silico* for potential dimers and hairpin structures at relevant thermal conditions using the Multiple Primer Analyzer (Thermo Fisher) and OligoAnalyzer (IDT).

We finally selected the primer pair MiAmphiL_28F (5’-CCTCGCCTGTTTACCAAAAAC**AYCGCCT**-3’) - MiAmphiL_extR (5’-**AAG**CTCCAT**R**GGGTCTTCTCGT**CTWRT**-3’) (Bases indicated in bold were added or modified from the primers of Sakata *et al*. 2022) to generate short-read amplicons (∼250 bp) from environmental samples. The primer pair MiAmphiL_28F - 16Sbr (5’-CCGGTCTGAACTCAGATCACGT-3’), combining the same forward primer with the ‘universal’ 16Sbr primer, was selected for obtaining long-read barcodes from skin swab samples collected from amphibian specimens. See Protocol S1 in the Supporting Information for additional recommendations on the design of RPA-compatible metabarcoding primers.

### RPA assay validation

We used the TwistAmp basic kit (TwistDX, Abbott) for RPA following the manufacturer’s instructions with deviations and two-step indexing for high-throughput sequencing as described in Plewnia et al. (2025b). In brief, the amphibian target fragment was amplified with a completely homologous version of the respective primers in a first RPA. In a subsequent RPA, these initial amplicons were amplified with a version of the same primer pair now carrying non-homologous 20-bp indexes as RPA’s only localized strand-displacement does not allow for index introduction in a single step. In addition, for environmental samples (in this case: eDNA water samples), we increased the template volume to 2.64 µl and did not add nuclease-free water to keep the reaction volume of 10 µl constant for the first RPA, while for the indexing RPA we used 1µl template only. RPA products were screened with gel electrophoresis for successful amplification.

### Library preparation and sequencing

Indexed RPA products were pooled at equal volumes and cleaned using magnetic beads (NucleoMag, Macherey-Nagel). We prepared libraries using the SQK-LSK114 kit following the manufacturer’s instructions for Flongle libraries. However, we changed the input volume, using 200 ng of DNA from the cleaned and combined pools as we previously experienced that the recommended 50-100 fmol led to high proportions of inactive pores when sequencing short amplicons. We further deviated cycling conditions for end-prep incubating for 30 min at 20°C and 30 min at 65°C. Sequencing was conducted for 20-30 h depending on pore activity on FLO-FLG114 Flongle flow cells using R10 chemistry and following the manufacturer’s instructions. For a detailed field-deployable protocol, please refer to Protocol S2 in the Supporting Information.

### Sequence processing and field-based reference assembly

We employed Dorado 0.9.0 for GPU-based basecalling of MinION reads using the super-accurate (sup) model. We then demultiplexed and trimmed sequence reads using minibar (Krehenwinkel et al. 2019) with default edit distances and filtered sequences for Phred scores >12 and read length of 172–212 bp for the eDNA marker and 520–580 bp for the barcoding marker using SeqKit (Shen et al. 2024). As we expected the community structure to be complex, including closely related species and unequal abundances for the eDNA marker, we refrained from OTU clustering approaches. Instead, we simply dereplicated identical sequences using Vsearch (Rognes et al. 2016) and mapped all unique sequences to a reference database following the direct BLASTn approach (Plewnia et al. 2025b). We chose the MIDORI2 reference database (MIDORI_UNIQ_NUC_SP_GB263_lrRNA_BLAST, based on a GenBank download from 13 October 2024; Leray et al. 2022), which is a readily available reference database for the long ribosomal subunit that contains all unique but no duplicated sequences from GenBank. With its small size, this database allows offline taxonomic assignment on a local machine without a need for internet connectivity or vast computational resources. For sequence assignment, we used BLASTn from blast+ 2.16.0 and subsequently filtered taxonomic hits as described in Plewnia et al. (2025b). As the MIDORI2 database already contains detailed taxonomic information, no subsequent ‘translation’ of accession numbers into species names was required.

For single-specimen barcoding, we built consensus sequences from filtered reads using NGSpeciesID (Sahlin et al. 2021), with the --ont option optimized for Nanopore reads and the medaka polishing algorithm (see Protocol S3 for details). Because NGSpeciesID may produce erroneous consensus sequences when target sequence abundance is low (pers. obs.), we visually examined all consensus sequences by aligning sequences using MAFFT with automated reverse complementing and subsequent polishing with CIAlign as described above. Consensus sequences were then manually trimmed in the alignment to remove divergent sequence ends that are sometimes erroneously introduced by NGSpeciesID. We subsequently inferred a maximum-likelihood phylogenetic tree using IQ-TREE 2 with 100,000 ultrafast bootstrap replicates and 100,000 replicates of the SH-like approximate likelihood ratio test (Minh et al. 2020), rooted with the caudate amphibian sequences generated (Table S2), to further validate the integrity of the curated consensus sequences (Figure S3).

Resulting sequences were stored as FASTA files with species name, unique identifier and sequence data stored in the header for incorporation in the reference database. Sequences were then concatenated into the FASTA-formatted version of the same MIDORI2 database and indexed into a searchable database using BLAST+. Subsequently, eDNA reads were blasted against the manually supplemented database as described above.

GenBank accession numbers for single-specimen 16S barcoding data generated in this paper are available in Table S2. Basecalled and demultiplexed raw MinION reads from both environmental samples and specimens are available in the supplementary data (Figshare). Associated scripts for building the integrated reference database and sequence processing are provided in Protocol S3 and Protocol S4 in the Supporting Information.

### Case study

To validate the performance of our eDNA assay, we collected water from 13 streams in the Sierra Nevada de Santa Marta of Colombia (see Figure S4; Table S3). This isolated mountain range is home to a diverse community of locally endemic amphibian species (Ruthven 1922; Pérez-Gonzalez et al. 2015; Rueda-Solano et al. 2016). Due to its remoteness and limited accessibility, the entire region remains under-researched and sequences for taxonomic reference are only available for a small subset of species (Arroyo et al. 2022; Lötters et al. 2025), which has so far impeded metagenetic biodiversity assessments. eDNA (water) samples were taken on site using a portable sampling device (Plewnia et al. 2025b). In addition to the eDNA samples, we collected non-invasive skin swabs from all encountered amphibian species at the same sampling sites, which were used to create a *de novo* reference database as described above

Samples were extracted using the DNeasy Blood and Tissue Kit (Qiagen) following the protocol of Plewnia et al. (2025a,b) for water filters and Böning et al. (2024) for amphibian swab samples. As the swab samples were also used for a parallel study on the same study system, we did not employ ‘fast-extraction’ techniques directly in the field. However, since we have shown previously that these techniques can yield sufficient RPA-ready template under field conditions for both amphibian skin swabs (Hoenig et al. 2023, 2024) and environmental samples (Plewnia et al. 2025b), our workflow remains entirely field-deployable (see Protocols S2-S4 in the Supporting Information). We classified species and genera as ‘incorrectly assigned’ from eDNA when they were neither visually observed and identified in our sampling sites during the present study, nor referenced in any previous works (Ruthven 1922; Pérez-Gonzalez et al. 2015; Rueda-Solano et al. 2016). Considering the near-complete assignment of eDNA reads to observed species using our *de novo* reference assembly (see results), this approach was deemed unlikely to falsely disregard undescribed amphibian diversity in our eDNA samples.

## Results

### An RPA and Nanopore-based workflow for field-deployable parallel species barcoding and eDNA metabarcoding

A schematic summary of the integrative workflow is presented in Figure 1. The workflow combines stepwise isothermal library preparation from environmental samples with non-invasive specimen barcoding for parallel on-site sequencing using portable Oxford Nanopore platforms. Consensus sequences from specimen barcoding can be used to supplement or build a site-specific reference database used for offline and field-deployable taxonomic assignment of sequences generated in parallel from environmental samples. RPA was able to amplify long fragments (∼600 bp) of 16S from amphibian skin swab samples, while short markers (∼250 bp) were most suitable for amplifying anuran eDNA from environmental samples. Please refer to Figure S2 for an overview of all primer tests. Complete field protocols and scripts used for sequence processing are provided in Protocols S2-S4 in the Supporting Information.

**Figure 1.**
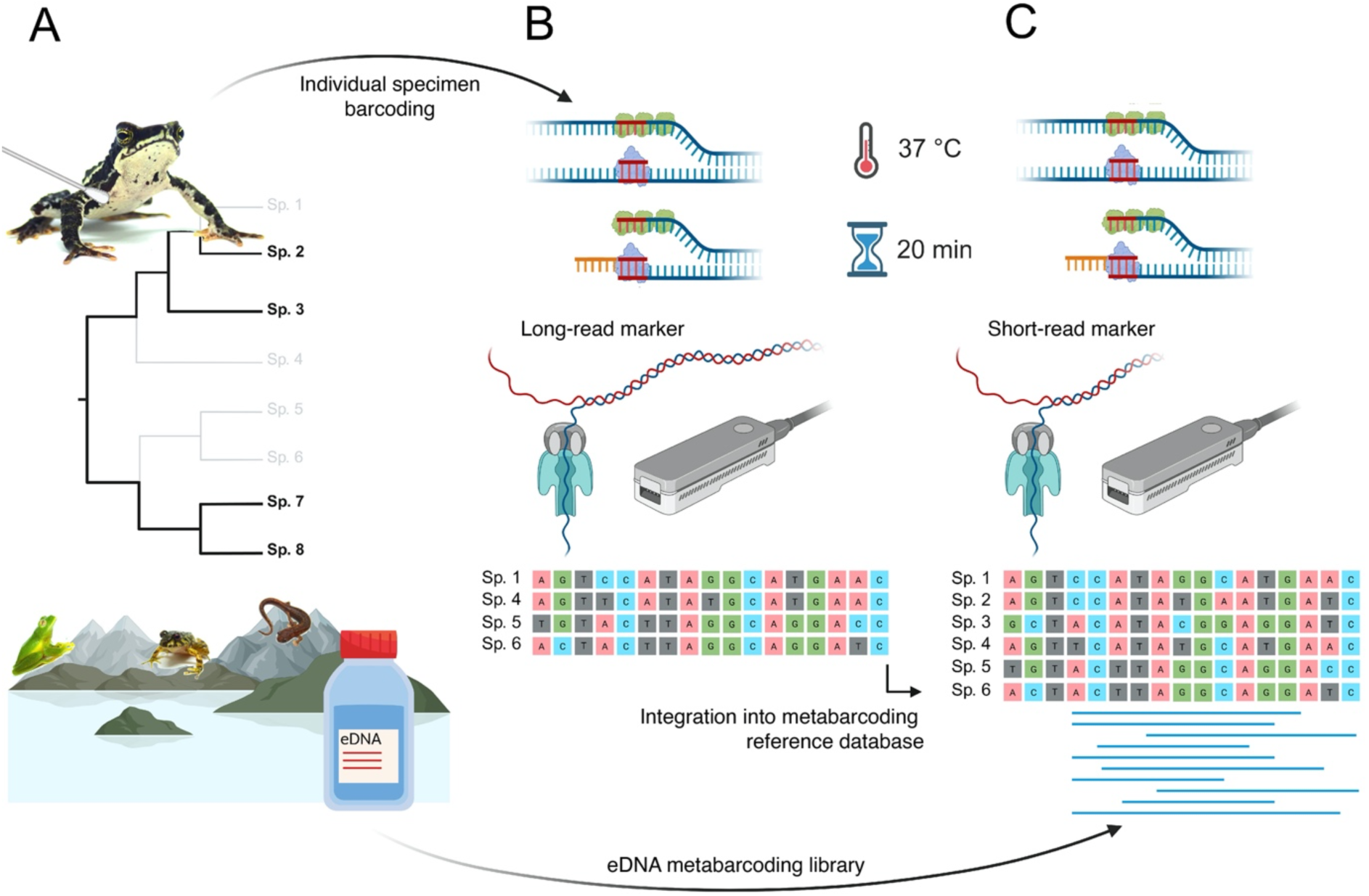
Overview of the novel RPA-based metabarcoding workflow with *de novo* assembly of a reference database using a two-step approach. (A) Environmental DNA (eDNA) sample collection is combined with non-invasive DNA sampling of target species that are absent or under-represented in existing reference datasets. Taxa theoretically lacking molecular reference data are displayed in grey in the phylogeny. (B) RPA-based amplification and indexing is combined with Oxford Nanopore MinION sequencing to generate long-read (∼600 bp) consensus sequences for barcoding individual specimens. (C) eDNA samples are prepared in parallel for high-throughput Oxford Nanopore MinION sequencing using a nested, short-read marker (∼250 bp). The resulting eDNA metabarcoding library is finally blasted to a reference database that is composed of a GenBank 16S rRNA subset supplemented with the consensus sequences generated in (B). In the RPA illustration, the green protein represents the single-stranded binding protein and the purple protein represents the recombinase.

### De novo reference database generation improves metabarcoding accuracy

In parallel with eDNA water sample collection, non-invasive DNA sampling of all amphibian morphospecies encountered around the same sites revealed the presence of 17 taxa based on 72 sampled individuals (Table S2; Figure S3). For ten of these taxa, comparative 16S sequence data were already available prior to our study (Figure S3). While we found that using prior available DNA reference data (MIDORI2; a mitochondrial rRNA GenBank subset) as a reference database for subsequent eDNA metabarcoding allowed a generalized taxonomic assignment at the genus level, *de novo* assembly of a combined, site-specific reference database (MIDORI2 integrated with the local single-specimen barcoding data from 72 sampled individuals) enabled a significantly improved taxonomic accuracy, most notably at the species level (Figure 2).

**Figure 2.**
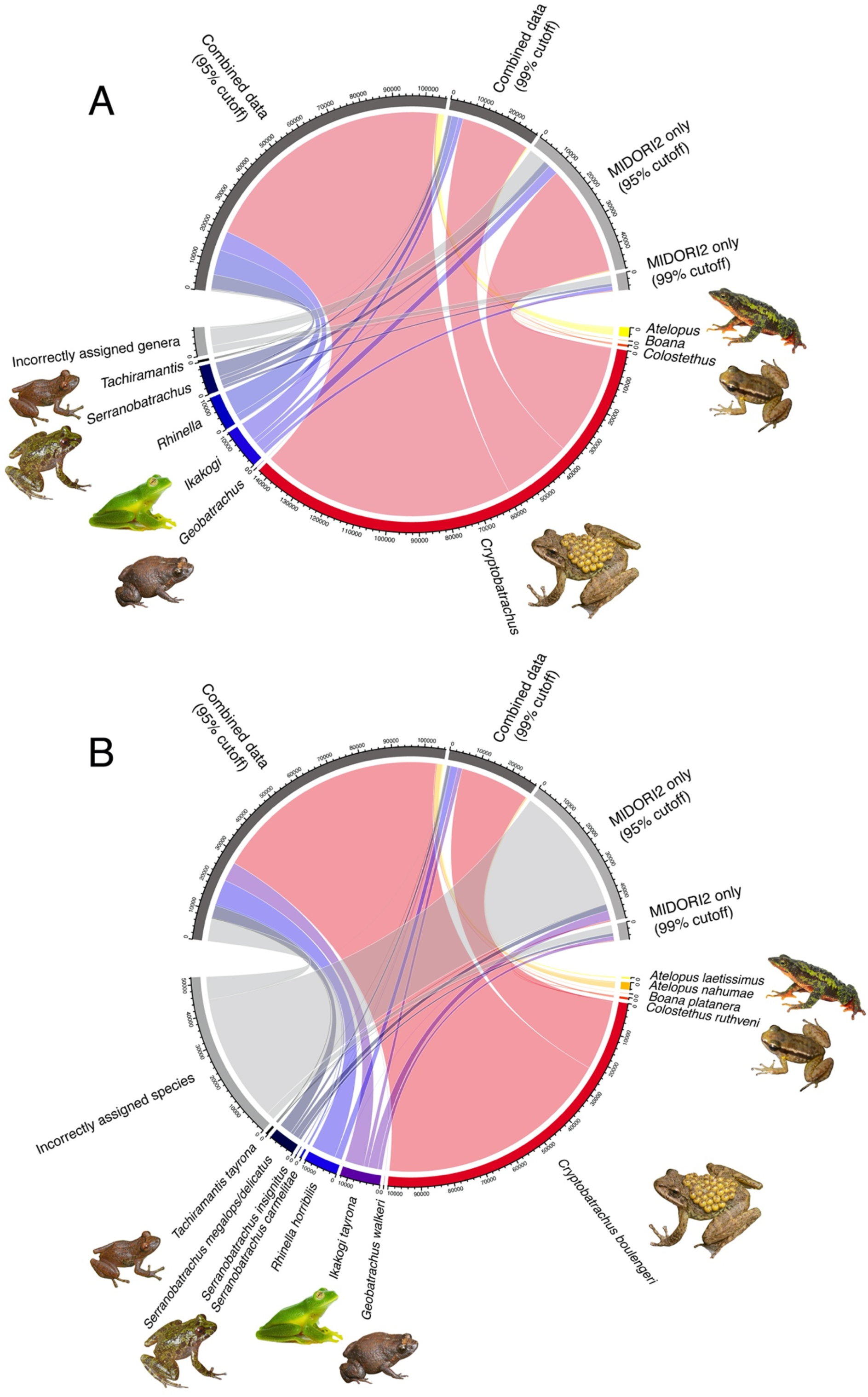
Summary of environmental DNA (eDNA) sequence reads, as obtained through isothermal library preparation and Nanopore sequencing and assigned to Operational Taxonomic Units (OTUs), based on 13 sampling sites within the Sierra Nevada de Santa Marta in northeastern Colombia. Incorrectly assigned species and genera were defined as those that were not visually observed and identified at the same sampling sites (Figure 3; Figure S3) nor referenced in any previous works. Two reference databases are compared, with the MIDORI2 mitochondrial GenBank subset being shown in comparison to the combined (*de novo* assembled) reference database (with local single-specimen barcodes integrated into the same MIDORI2 dataset). Comparisons use both a 95% and 99% sequence identity (BLASTn) cutoff for accepting the assigned taxonomy, with results summarized on (A) genus and (B) species level. This comparison includes only the amphibian taxa recovered above the respective sequence similarity threshold, with the numbers corresponding to the number of amphibian sequence reads for each database format.

At the 95% sequence similarity cutoff, amphibian reads constituted 61.8% (total: 53,225) of all reads using the combined dataset and 59.4% (total: 48,091) of all reads using the MIDORI2-only dataset. The combined dataset yielded 99.5% correct genus-level and 93.1% species-level assignment of amphibian reads, compared to 88.3% and 12.2% for the MIDORI2-only dataset, respectively. This corresponds to a 13% higher probability of correct read assignment at the genus level (RR = 1.13, 95% CI: 1.12–1.31, *p* < 0.001) but a more than seven times greater likelihood of correct species-level assignment (RR = 7.63, 95% CI: 7.45–7.82, *p* < 0.001) when using the combined dataset. At this cutoff, the MIDORI2-only dataset identified 6 visually confirmed species, whereas the combined dataset identified 12 species.

At the more stringent 99% sequence similarity cutoff, amphibian reads constituted 68.9% (total: 28,302) of all reads using the combined dataset and 29.4% (total: 5,329) of all reads using the MIDORI2-only dataset. The combined dataset showed a near-complete correct assignment of all amphibian reads (99.8%), with all correct genus-level reads also resolving to the species level. The MIDORI2-only dataset showed a substantially lower taxonomic accuracy (genus: 50.9%; species: 50.7%), meaning the probability of correct genus-and species-level assignment was around twice as high when using the combined dataset (genus: RR = 1.96, 95% CI: 1.91–2.01, *p* < 0.001; species: RR = 1.97, 95% CI: 1.92–2.02; *p* < 0.001). At this cutoff, the MIDORI2-only dataset identified 5 visually confirmed species, while the combined dataset identified 10 species.

Non-amphibian OTUs of other locally known taxa largely corresponded to other vertebrates, with reads most commonly assigned to common opossum (*Didelphis marsupialis*), catfish (*Trichomycterus* spp., *Cordylancistrus tayrona*), the Colombian red howler monkey (*Alouatta seniculus*), and cattle (*Bos taurus*; in the lower regions). For an overview of total read counts and taxon assignments across the different field sites and reference database formats, please refer to the Supporting Data.

### De novo reference database construction allows real-time characterization of community structure

We employed the novel assay across 13 eDNA sampling localities situated at various elevations in the Sierra Nevada de Santa Marta mountain range in northeast Colombia. By integrating local single-specimen barcoding data into the reference database prior to eDNA analyses, a total of 12 visually confirmed amphibian species could be recovered in these environmental samples (using a 95% sequence similarity cutoff; Figure 2B). The inclusion of field and RPA replicates (Table S3) notably increased species detection probabilities across localities, with a mean 1.74 amphibian species detected per indexed sample, but a mean 2.92 per field site.

Our eDNA results confirmed an elevational structure in species distributions, with the lowland generalist species *Rhinella horribilis* and *Boana platanera* found only in the lowest site, and the montane endemics *Atelopus laetissimus, Geobatrachus walkeri*, and *Serranobatrachus carmelitae* found only in samples above 2,000 m elevation (Figure 3). Species detection from eDNA was more consistent for species with aquatic larval stages (the genera *Atelopus, Boana, Colostethus, Ikakogi*, and *Rhinella*) or stream-associated occurrence (*Cryptobatrachus boulengeri* and *S. carmelitae*), showing no large spatial gaps between sampling sites in their elevational range. These patterns were reflected by absolute read counts, with stream-breeding and associated species presenting the vast majority of assigned reads from water eDNA samples across our study sites (Figure 2B; Figure 3). Instead, detection of terrestrial or arboreal forest-associated direct-developers [the genera *Geobatrachus*, *Serranobatrachus* (except *S. carmelitae*), and *Tachiramantis*] was more inconsistent considering the elevational range in which they have directly been observed (Figure 3; Table S4). Four anuran species (*Serranobatrachus sanctaemartae*, *S. cristinae*, *Lithobates vaillanti*, and an undescribed *Tachiramantis* sp.) that were observed directly and included in single-specimen barcoding were not recovered in any of the eDNA samples. With the exception of *L. vaillanti*, these taxa are similarly associated with a terrestrial and/or arboreal life history (Table S4).

**Figure 3.**
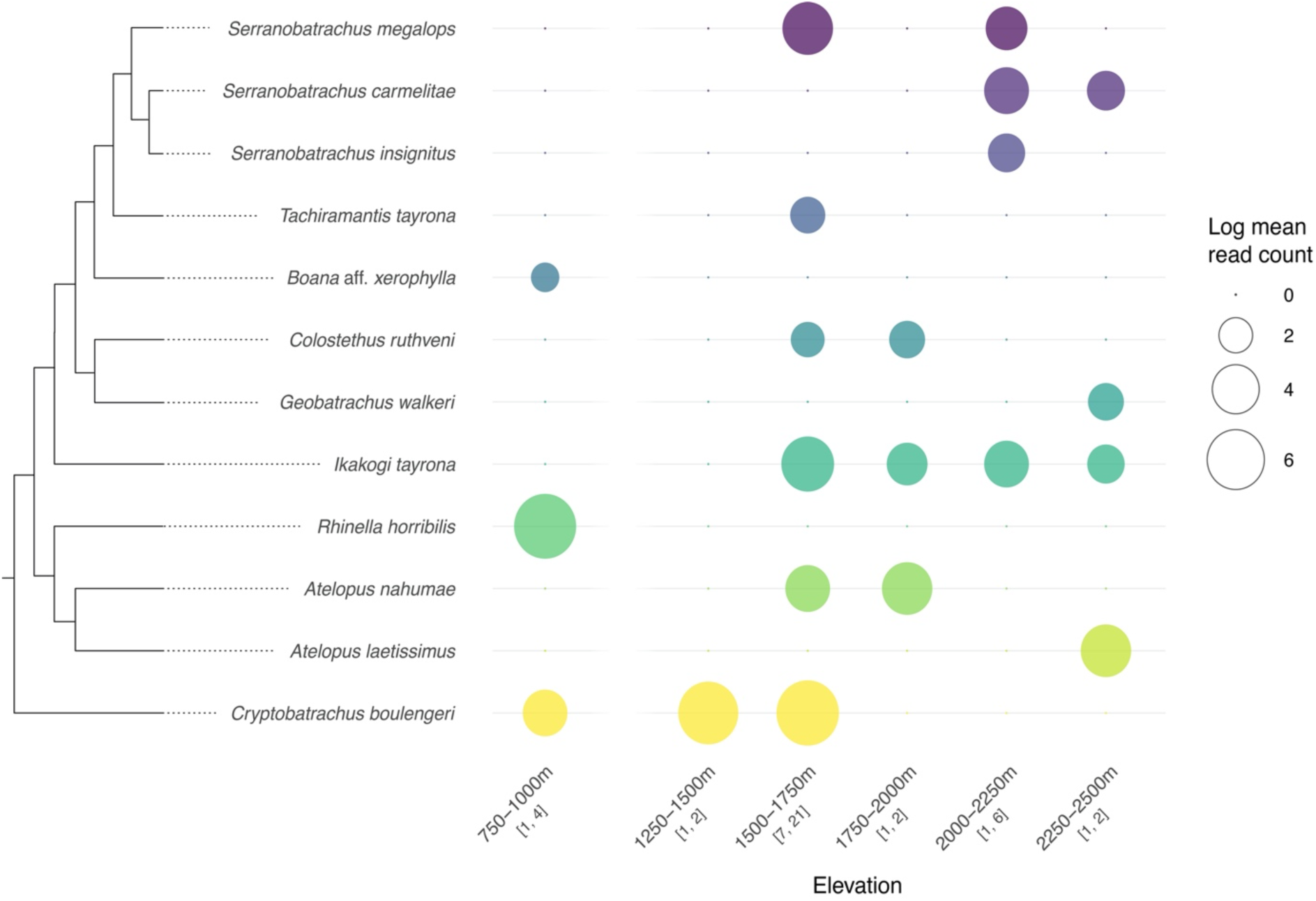
Environmental DNA (eDNA) sequencing read counts for all species that were correctly assigned to species (*i*.*e*., species observed directly and identified, see Figure 2B) across elevations in the Sierra Nevada de Santa Marta in northeastern Colombia. The data are presented as the natural logarithm of the mean read counts. The number of sampling sites (first value) and total sequenced field and RPA replicates (second value) per elevational category are indicated in brackets. Note that no sampling sites were situated between 1,000-1,250 m, and eDNA sampling comprised a non-homogenous spread across elevations. See Table S3 for details on field sites and replicates. Phylogenetic relationships display those inferred from single-specimen barcoding (refer to Figure S3 for a Maximum Likelihood tree of all barcoded species and support values).

## Discussion

With precipitous and ongoing global biodiversity declines presenting one of the most urgent challenges of our time, leveraging the use of molecular species monitoring tools becomes increasingly important to enable timely conservation action (Altermatt et al. 2025). We here introduce a simplified RPA-and Nanopore MinION-based workflow that uses non-invasive barcoding of species lacking molecular reference data, and integrate these with existing sequence data to create a highly accurate reference database for community metabarcoding for direct use in the field. Focusing on a threatened and remote tropical amphibian assemblage in northern Colombia, we demonstrated that *de novo* reference assembly strongly increased the taxonomic accuracy of metabarcoding assessments, doubled the number of visually confirmed species detected in eDNA samples, and increased the proportion of correct species-level assignments of eDNA sequence reads from 12.2% to 93.1% at a 95% sequence identity cutoff. At a more stringent 99% sequence similarity cutoff, correct species-level assignment rose from ∼50% to near-complete accuracy. In turn, employing our two-step barcoding assay enabled a better characterization of the altitudinal community structure of amphibian species. However, as commonly observed for eDNA from water samples, detection was more consistent for species with aquatic larvae or stream-associated life histories than for arboreal or terrestrial species. Several, mostly terrestrial species present in the combined (*de novo* assembled) reference database were not detected in eDNA samples, reflecting the life history of species with a reduced likelihood of shedding genetic material into aquatic systems (de Souza et al. 2016, Ruppert et al. 2019, Davis et al. 2025).

*De novo* reference sequence generation not only enhanced resolution but also strongly increased the number of reads passing the similarity thresholds and being assigned to target taxa, implying that a lower sequencing depth may be sufficient for confident assessments. This is a critical consideration when operating with limited flow cell capacity (such as the Flongle flow cell used in this study) and time constraints in remote field settings (De la Cerda et al. 2023, *cf*. Zorz et al. 2023). Further advances in taxonomic resolution could be achieved by targeting longer markers for metabarcoding, as increasingly feasible with third-generation sequencing (Heeger et al., 2018; Jamy et al., 2020; Ohta et al., 2023; Ip et al. 2025; Plewnia et al. 2025b). For environmental samples, however, marker length is constrained by DNA quality (Brandão-Dias et al. 2025), the likely cause for only short markers consistently amplifying environmental samples with RPA in our dataset. Indeed, long markers may ‘overlook’ rare species in environmental samples where DNA fragmentation prevents amplification (Doorenspleet et al. 2025). A solution for this could be the use of several, stepwise nested markers, where complete community composition is approximated by a short marker, nested within a longer target fragment that allows increased taxonomic resolution for more abundant species where detection of less degraded DNA (*i.e.* allowing long-read amplification) is more likely (Stoeckle et al. 2018, Ruppert et al. 2019, Brandão-Dias et al. 2025). Our final primer selection represents a fraction of all possible combinations and, in addition, we could not exhaustively test the functionality of all primer pairs. Optimizing and scaling our approach will strongly benefit from a better automated metabarcoding primer design pipeline that takes RPA-specific requirements such as longer primers and higher mismatch tolerance into account. Several of the RPA primers we designed and utilized for our study successfully amplified DNA from a broad range of amphibian taxa from environmental samples. Furthermore, the primers yielded partial metabarcoding ‘bycatch’, mostly in the form of other vertebrate taxa, which can add valuable information for exhaustive habitat-scale biodiversity assessments (Plewnia et al. 2025a).

Our workflow can be readily adapted to other taxonomic groups and ecosystems where reference coverage remains poor, thus directly contributing to the establishment of comparative molecular data for regions that are particularly understudied (Marques et al. 2021). Future developments for the field-deployable eDNA toolkit may explore amplification-free approaches such as isothermal and non-invasive CRISPR-capture strategies (Plewnia et al. 2025c). Coupled with MinION sequencing, this could allow simultaneous species detection and methylation profiling from ‘native’ DNA. Such methods hold promise for adding demographic and health indicators to biodiversity assessments (e.g. Crossman et al. 2021; Zhao et al. 2023; Le Clercq et al. 2023; Anderson et al. 2024). Further improvements to field-based metabarcoding assays may include affinity-tagged primers which, in combination with lateral flow detection, could allow visualization of amplification success prior to pooling and sequencing on site with minimal additional equipment (Lobato and O’Sullivan, 2018, Plewnia et al. 2025c).

In summary, by integrating single-specimen reference generation directly into a PCR-free field-deployable metabarcoding workflow, we can bridge critical data gaps that are currently impeding accurate molecular biodiversity monitoring in high-diversity but understudied ecosystems (Ip et al. 2025). Our approach is portable, rapid, and scalable, providing practitioners in remote and resource-limited settings with the capacity to generate high-quality, community-level biodiversity data, with environmental sampling and species-specific sampling being carried out in parallel. Applying this workflow in biodiversity hotspots can accelerate the creation of essential baseline ecological data, and strengthen decision-making for conservation in the very regions where timely action is most urgently needed.

## Supplementary Material

All data underlying this paper are provided in the Supporting Information, and additional supplementary data (demultiplexed MinION raw reads and read count tables) can be found on the Figshare open-access repository at www.doi.org/10.6084/m9.figshare.28417451.

## Acknowledgments.

We are grateful to Philipp Böning, Till-Hendrik Macher and Yannis Schöneberg for valuable discussion, to Karin Fischer, Susan Kennedy and Sabine Naber for labwork support, and to Aldair Barros, Jose D. Barros Castañeda, Romario Salas and Tobias Hildwein for help in the field. Stephan Seeling helped with funding acquisition. This work was partially funded under the project Biodiversitätsmonitoring 2.0 by Ministerium für Wirtschaft, Verkehr, Landwirtschaft und Weinbau Rheinland-Pfalz. Sample collection was partially funded by the Deutsche Gesellschaft für Herpetologie und Terrarienkunde and the Forschungsfonds of Trier University. Samples used in this study were collected and exported under permits P02182S9571_N0055 and Permiso no CITES 003790 granted by Autoridad Nacional de Licencias Ambientales, ANLA, Colombia.

## Author Contributions

JE, AP and CH initiated the idea and conceptualized the study. JE conducted lab work with contributions from AP and CH. JE, CH and AP analysed and visualized the data. AJC and LAR-S helped with sample and permit acquisition. SL and HK provided project resources and administration. JE and AP wrote the first draft. All authors revised and edited the manuscript.

