## Supporting Information for "A field-deployable eDNA metabarcoding workflow including *de novo* reference assembly for characterizing understudied biodiversity hotspots"

**Table S1**. List of primers tested for use with Recombinase Polymerase Amplification (RPA). Bases that were edited or added with respect to previously published primers (see references) are highlighted in bold. Primers not adapted from existing references were designed by visually scanning the reference alignment (Portik *et al.* 2023) for conserved regions. For the column 'Site': numbers and colours correspond to the different primer binding sites on the 16S rRNA gene as illustrated in Figure S1. Dots highlight the primer pairs selected for use in final analyses (• Long-read marker for DNA isolates; •• Short-read marker for eDNA samples; also see Table S2).

| Primer name | F/R | Sequence (5'-3') | Reference | Site |
|---|---|---|---|---|
| **MiAmphiL_ext28_F ••** | F | CCTCGCCTGTTTACCAAAAAC**AYCGCCT** | Sakata *et al.* 2022 | 1 |
| MiAmphiL_ext30_F | F | **R**CCTCGCCTGTTTACCAAAAAC**AYCGCCTY** | | |
| MiA16SF_25ext | F | GCCTGTTTACCAAAAAC**AYCGCCT** | | |
| MiA16SRasF_27ext1 | F | **AYWAG**ACGAGAAGACCCCATGGAG**CTT** | Sakata *et al.* 2022 (modified: reverse as forward) | 2 |
| MiA16SRasF_27ext2 | F | **AYAAG**ACGAGAAGACCCCATGGAG**CTT** | | |
| MiA16SRasF_25ext | F | **AAG**ACGAGAAGACCCCATGGAG**CTT** | | |
| MiA16SRasF_25extD | F | **AAG**ACGAGAAGACCC**Y**ATGGAG**CTT** | | |
| **MiAmphiL_ext_R •** | R | **AAG**CTCCAT**R**GGGTCTTCTCGT**CTWRT** | Sakata *et al.* 2022 | 3 |
| MiA16SR_27ext1 | R | **AAG**CTCCATGGGGTCTTCTCGT**CTWRT** | | |
| MiA16SR_27ext2 | R | **AAG**CTCCATGGGGTCTTCTCGT**CTTRT** | | |
| MiA16SR_25ext | R | **AAG**CTCCATGGGGTCTTCTCGT**CTT** | | |
| MiA16SR_25extD | R | **AAG**CTCCAT**R**GGGTCTTCTCGT**CTT** | | |
| 16S_APJE_R1 | R | GCGCTGTTATCCCYAGGGTAACTTGGTTC | No reference used | 4 |
| 16S_APJE_R2 | R | GCTGTTATCCCYAGGGTAACTTGGTTC | | |
| 16S_APJE_R3 | R | GCGCTGTTATCCCYAGGGTAACTTGGT | | |
| 16S_APJE_R4 | R | TGTTATCCCYAGGGTAACTTGGTTC | | |
| 16S_preMig_R1 | R | CCTGATCCAACATCGAGGTCGTAAACC | No reference used | 5 |
| 16S_preMig_R2 | R | TGATCCAACATCGAGGTCGTAAACC | | |
| 16S_preMig_R3 | R | CCTGATCCAACATCGAGGTCGTAAACC | | |
| 16S_preMig_R4 | R | CCTGATCCAACATCGAGGTCGTAAACC | | |
| **16SbH •** | R | CCGGTCTGAACTCAGATCACGT | Palumbi *et al.* 2002 | 6 |
| 16SbH_ext | R | **CY**CCGGTCTGAACTCAGATCACGT**RRGGYWT** | | |
| 16SbH_ext_28 | R | **CY**CCGGTCTGAACTCAGATCACGT**RRGG** | | |
| 16SbH_ext_26 | R | CCGGTCTGAACTCAGATCACGT**RRGG** | | |
| 16SbH_ext_26nd | R | CCGGTCTGAACTCAGATCACGT**AGGG** | | |

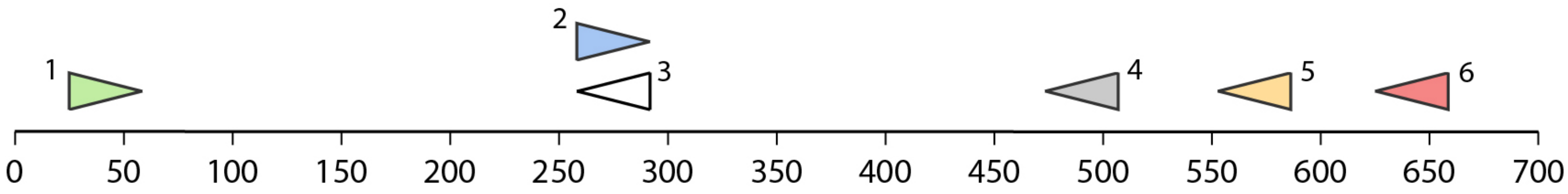


**Figure S1**. Map of primer binding sites on the 16S rRNA mitochondrial gene. Numbers and colours correspond to the primer overview in Table S1. The positions are in reference to *Xenopus laevis* in the 16S alignment by Portik *et al.* (2023). Note that positions may vary between species.

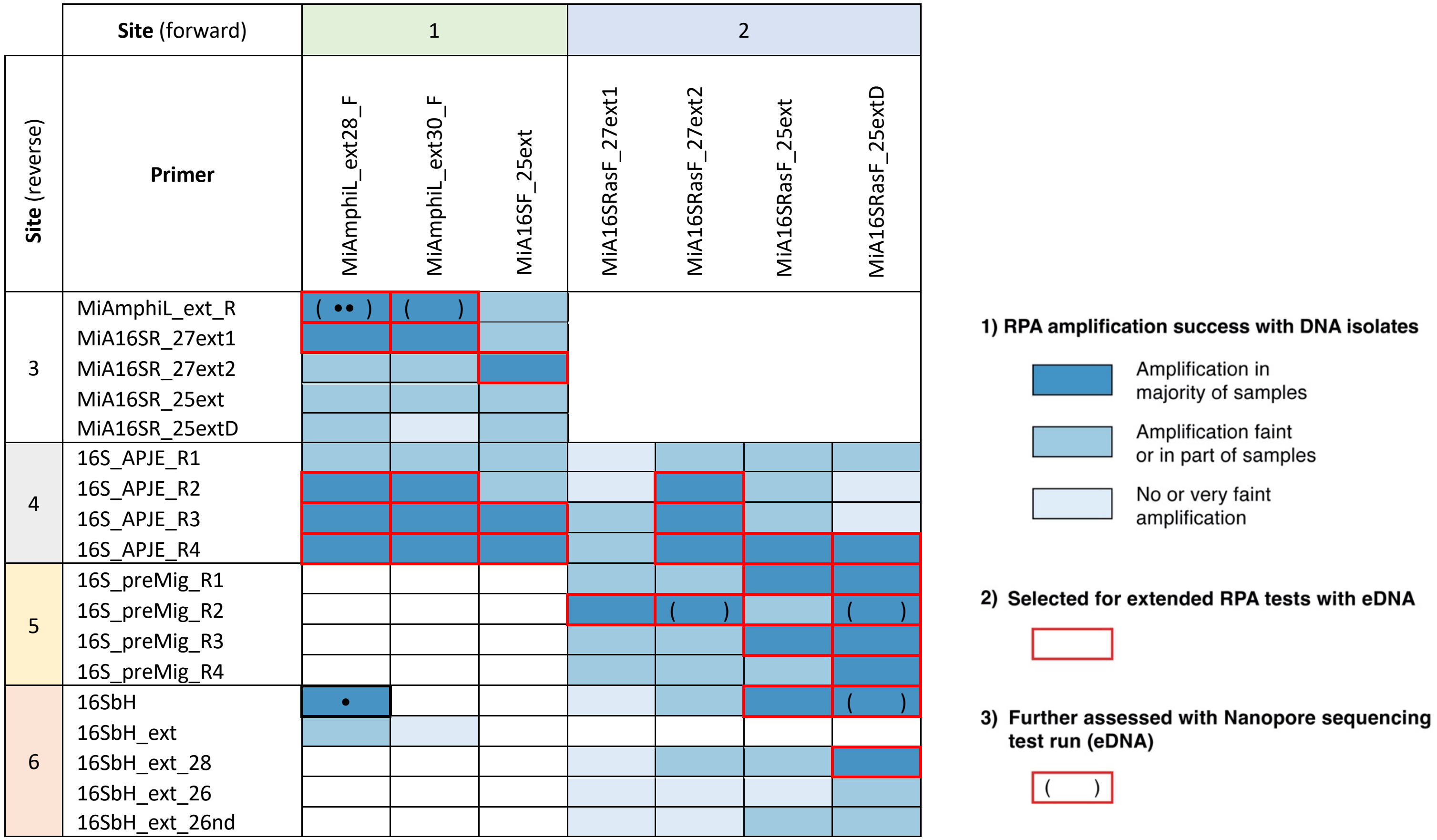


**Figure S2**. Overview of primer combinations tested for use with Recombinase Polymerase Amplification (RPA). Following RPA tests on DNA isolates from non-invasive amphibian samples (skin swabs), primer combinations showing successful amplification (based on visualization with gel electrophoresis; see legend) were taken for extended testing with environmental DNA (eDNA) samples (outlined in red). Given resource limitations, not all primer combinations could be tested on a broader subset of samples - primer combinations in white could not be tested or only on a single sample, precluding an assessment of their functionality. For eDNA samples, final primer selection was based on a Nanopore sequencing test run. Dots highlight the primer pairs selected for use in final analyses (• Long-read marker for DNA isolates; •• Short-read marker for eDNA samples). For the primer binding sites, numbers and colours correspond to Table S1 and Figure S1.

**Table S2.** GenBank accession numbers for 16S sequences generated from non-invasive skin swabs of amphibian species in the Sierra Nevada de Santa Marta, northern Colombia. Only generalized locality names are provided to protect already threatened populations (Lindenmayer & Scheele, 2017).

| Species | Locality | GenBank # |
|---|---|---|
| *Bolitoglossa savagei* | Sogrome | PX105574 |
| *Bolitoglossa savagei* | San Pedro | PX105575 |
| *Bolitoglossa savagei* | San Pedro | PX105576 |
| *Serranobatrachus megalops/delicatus* | Cuchillo San Lorenzo | PX105577 |
| *Serranobatrachus megalops/delicatus* | Cuchillo San Lorenzo | PX105578 |
| *Serranobatrachus megalops/delicatus* | Cuchillo San Lorenzo | PX105579 |
| *Serranobatrachus megalops/delicatus* | Cuchillo San Lorenzo | PX105580 |
| *Serranobatrachus megalops/delicatus* | Cuchillo San Lorenzo | PX105581 |
| *Serranobatrachus megalops/delicatus* | San Pedro | PX105582 |
| *Serranobatrachus megalops/delicatus* | San Pedro | PX105583 |
| *Serranobatrachus sanctaemartae* | Cuchillo San Lorenzo | PX105584 |
| *Serranobatrachus carmelitae* | Cuchillo San Lorenzo | PX105585 |
| *Serranobatrachus sanctaemartae* | San Pedro | PX105586 |
| *Serranobatrachus cristinae* | Cuchillo San Lorenzo | PX105587 |
| *Serranobatrachus cristinae* | Cuchillo San Lorenzo | PX105588 |
| *Serranobatrachus cristinae* | Cuchillo San Lorenzo | PX105589 |
| *Serranobatrachus cristinae* | Cuchillo San Lorenzo | PX105590 |
| *Serranobatrachus insignitus* | San Pedro | PX105591 |
| *Serranobatrachus insignitus* | Cuchillo San Lorenzo | PX105592 |
| *Serranobatrachus insignitus* | San Pedro | PX105593 |
| *Serranobatrachus* cf. *carmelitae* | San Pedro | PX105594 |
| *Serranobatrachus* cf. *carmelitae* | San Pedro | PX105595 |
| *Serranobatrachus carmelitae* | San Pedro | PX105596 |
| *Serranobatrachus carmelitae* | San Pedro | PX105597 |
| *Tachiramantis tayrona* | Cuchillo San Lorenzo | PX105598 |
| *Tachiramantis tayrona* | Cuchillo San Lorenzo | PX105599 |
| *Tachiramantis* sp. | Río Tucurinca | PX105600 |
| *Tachiramantis* sp. | Río Tucurinca | PX105601 |
| *Tachiramantis* sp. | Cuchillo San Lorenzo | PX105602 |
| *Tachiramantis* sp. | Río Tucurinca | PX105603 |
| *Tachiramantis* sp. | Río Tucurinca | PX105604 |
| *Tachiramantis* sp. | Cuchillo San Lorenzo | PX105605 |
| *Tachiramantis* sp. | Río Tucurinca | PX105606 |
| *Tachiramantis* sp. | Cuchillo San Lorenzo | PX105607 |
| *Tachiramantis* sp. | Cuchillo San Lorenzo | PX105608 |
| *Tachiramantis* sp. | Cuchillo San Lorenzo | PX105609 |
| *Lithobates vaillanti* | Palmor | PX105610 |
| *Lithobates vaillanti* | Palmor | PX105611 |
| *Colostethus ruthveni* | Palmor | PX105612 |
| *Colostethus ruthveni* | Río Tucurinca | PX105613 |
| *Colostethus ruthveni* | Cuchillo San Lorenzo | PX105614 |
| *Boana* aff. *xerophylla* | Palmor | PX105615 |
| *Boana* aff. *xerophylla* | Palmor | PX105616 |
| *Boana* aff. *xerophylla* | Palmor | PX105617 |
| *Geobatrachus walkeri* | Cuchillo San Lorenzo | PX105618 |
| *Geobatrachus walkeri* | Cuchillo San Lorenzo | PX105619 |
| *Cryptobatrachus boulengeri* | Cuchillo San Lorenzo | PX105620 |
| *Cryptobatrachus boulengeri* | Cuchillo San Lorenzo | PX105621 |
| *Cryptobatrachus boulengeri* | Cuchillo San Lorenzo | PX105622 |
| *Cryptobatrachus boulengeri* | Cuchillo San Lorenzo | PX105623 |
| *Cryptobatrachus boulengeri* | Cuchillo San Lorenzo | PX105624 |
| *Cryptobatrachus boulengeri* | San Pedro | PX105625 |
| *Cryptobatrachus boulengeri* | Palmor | PX105626 |
| *Cryptobatrachus boulengeri* | Río Tucurinca | PX105627 |
| *Cryptobatrachus boulengeri* | Río Tucurinca | PX105628 |
| *Ikakogi tayrona* | Cuchillo San Lorenzo | PX105629 |
| *Ikakogi tayrona* | Cuchillo San Lorenzo | PX105630 |

| | | |
|---|---|---|
| *Ikakogi tayrona* | Cuchillo San Lorenzo | PX105631 |
| *Rhinella horribilis* | Palmor | PX105632 |
| *Rhinella horribilis* | Palmor | PX105633 |
| *Rhinella horribilis* | Palmor | PX105634 |
| *Atelopus nahumae* | Cuchillo San Lorenzo | PX105635 |
| *Atelopus nahumae* | Cuchillo San Lorenzo | PX105636 |
| *Atelopus nahumae* | Cuchillo San Lorenzo | PX105637 |
| *Atelopus nahumae* | Cuchillo San Lorenzo | PX105638 |
| *Atelopus nahumae* | Cuchillo San Lorenzo | PX105639 |
| *Atelopus laetissimus* | San Pedro | PX105640 |
| *Atelopus laetissimus* | Cuchillo San Lorenzo | PX105641 |
| *Atelopus laetissimus* | Cuchillo San Lorenzo | PX105642 |
| *Atelopus laetissimus* | Cuchillo San Lorenzo | PX105643 |
| *Atelopus laetissimus* | Cuchillo San Lorenzo | PX105644 |
| *Atelopus laetissimus* | Cuchillo San Lorenzo | PX105645 |

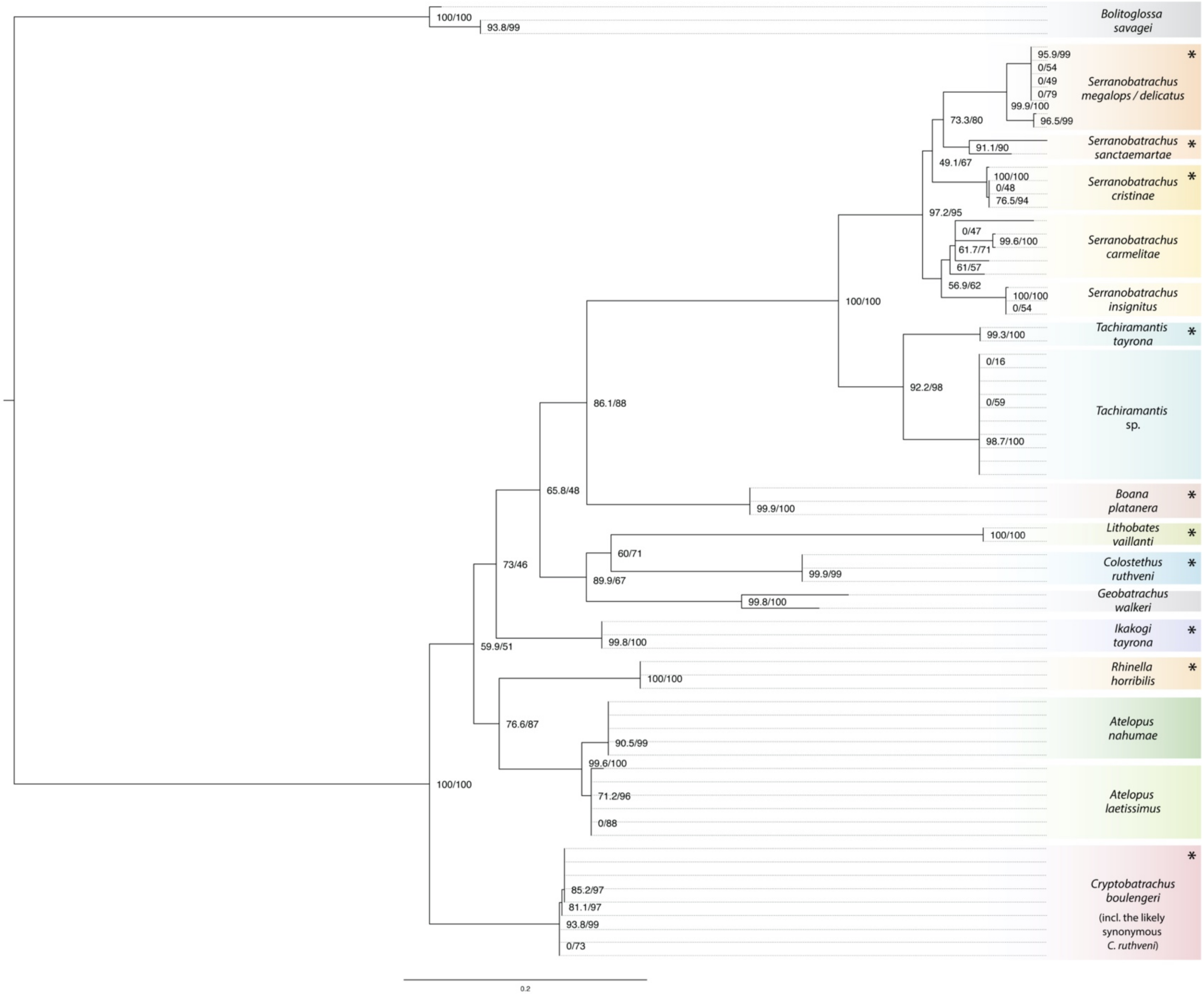


**Figure S3.** Maximum Likelihood tree inferred from a long-read 16S rRNA alignment (559 bp) obtained using Recombinase Polymerase Amplification (RPA) and Nanopore sequencing of non-invasive skin swab samples of species encountered in our study system, the Sierra Nevada de Santa Marta in northern Colombia. Node values designate ultrafast bootstrap and approximate likelihood ratio test (aLRT) values, respectively, based on 100.000 pseudoreplicates. Representing 17 species-level lineages (Of which 16 are anuran species, *Bolitoglossa* was used as outgroup), these single-specimen sequencing data were integrated with the MIDORI2 mitochondrial Genbank subset to constitute a *de novo* reference dataset to use in environmental metabarcoding using a nested, short-read marker. An asterisk denotes taxa for which prior 16S sequences were available in the MIDORI2 (GenBank subsetted) reference database prior to our study. *Boana platanera* is a recently described species that was previously included as *Boana* cf. *xerophylla* in GenBank and we considered a ‘correct’ species assignment accordingly.

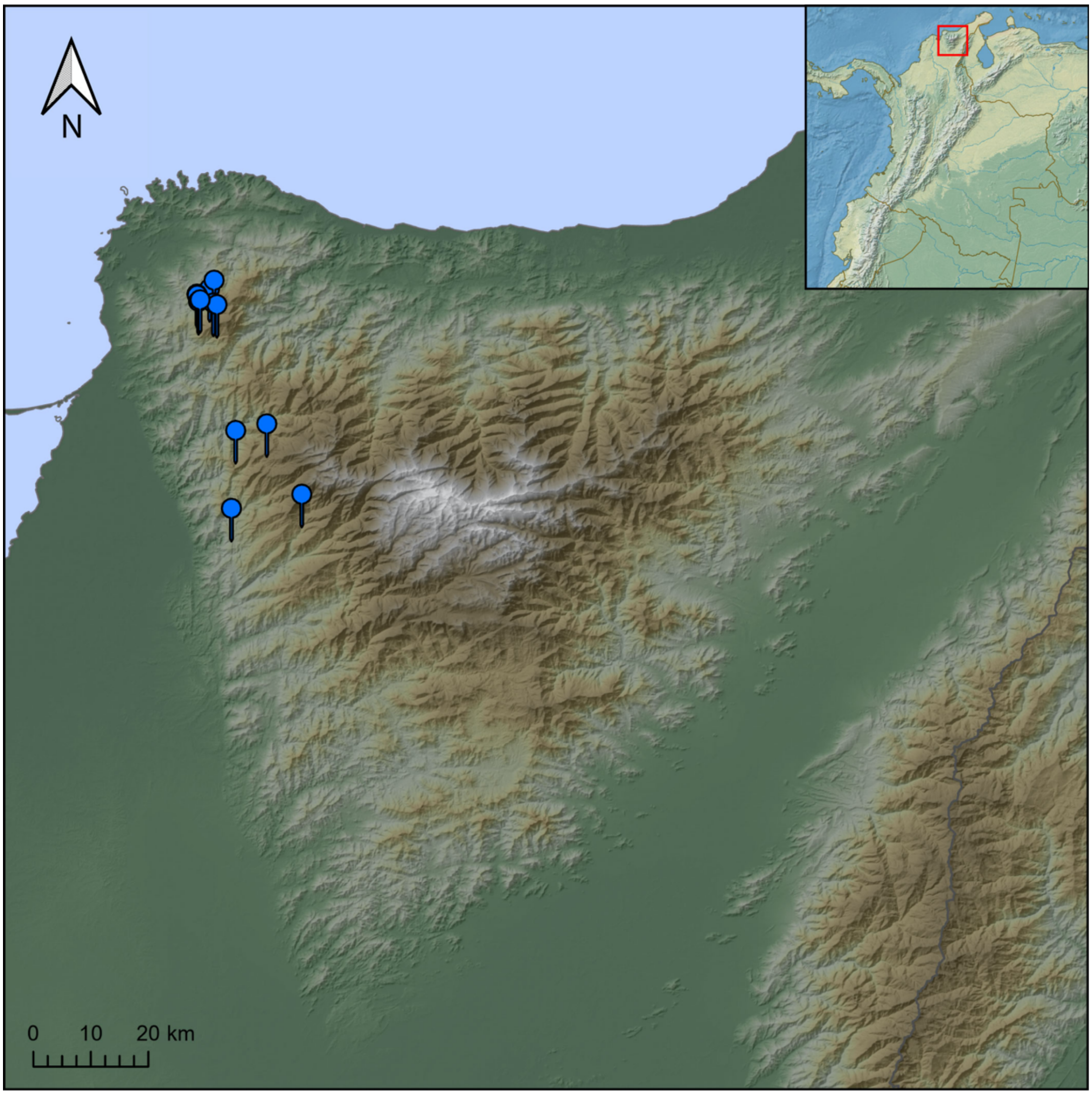


**Figure S4.** Map of eDNA sampling sites in the Sierra Nevada de Santa Marta (n = 13), with the inset showing its position within Colombia. Basemap adapted from Esri. Inset adapted from Natural Earth (naturalearthdata.com).

**Table S3.** Overview of eDNA sampling sites with the total number of field sampling and RPA replicates as included in the final Nanopore (MinION) sequencing run. For site 13, no accurate elevation is available.

| Site | Elevation | Locality | eDNA samples | RPA replicates | Total sequenced |
|---|---|---|---|---|---|
| 1 | 967 | Palmor | 2 | 2 | 4 |
| 2 | 1486 | Cuchillo San Lorenzo | 1 | 2 | 2 |
| 3 | 1505 | Cuchillo San Lorenzo | 1 | 2 | 2 |
| 4 | 1524 | Cuchillo San Lorenzo | 1 | 2 | 2 |
| 5 | 1542 | Cuchillo San Lorenzo | 1 | 2 | 2 |
| 6 | 1550 | Cuchillo San Lorenzo | 1 | 2 | 2 |
| 7 | 1557 | San Pedro | 2 | 1 + 2 | 3 |
| 8 | 1577 | Río Tucurinca | 2 | 2 | 4 |
| 9 | 1667 | Cuchillo San Lorenzo | 3 | 2 | 6 |
| 10 | 1766 | Cuchillo San Lorenzo | 1 | 2 | 2 |
| 11 | 2111 | Cuchillo San Lorenzo | 3 | 2 | 6 |
| 12 | 2459 | San Pedro | 1 | 2 | 2 |
| 13 | NA | Cuchillo San Lorenzo | 1 | 2 | 2 |

**Table S4.** Anuran species visually observed in the Sierra Nevada de Santa Marta (SNSM) around each respective eDNA sampling locality, including present knowledge on species endemism, habitat characteristics and reproductive mode. Question marks indicate likely presence on the basis of acoustic observations or by proximity to known records. For site 13, no accurate elevation is available.

| | Site | | | | | | | | | | | | | | |
|---|---|---|---|---|---|---|---|---|---|---|---|---|---|---|---|
| **Species** | **1** | **2** | **3** | **4** | **5** | **6** | **7** | **8** | **9** | **10** | **11** | **12** | **13** | **Endemism** | **Habitat type** | **Reproduction** |
| *Atelopus nahumae* | N | N | N | N | N | N | N | Y | N | Y | Y | N | N | SNSM endemic | Riparian (vegetation) | Aquatic larvae (lotic) |
| *Atelopus laetissimus* | N | N | N | N | N | N | N | N | N | N | Y | Y | N | | Riparian (vegetation) | Aquatic larvae (lotic) |
| *Ikakogi tayrona* | N | Y | Y | Y | Y | Y | Y | Y | Y | Y | Y | Y | Y | | Riparian (vegetation) | Aquatic larvae (lotic) |
| *Colostethus ruthveni* | Y | ? | ? | ? | ? | ? | Y | Y | ? | ? | N | N | ? | | Terrestrial | Aquatic larvae (lentic) |
| *Cryptobatrachus boulengeri* (incl. the likely synonymous *C. ruthveni*) | Y | Y | Y | Y | Y | Y | Y | Y | Y | Y | N | N | Y | | Riparian (boulders) | Direct development |
| *Tachiramantis tayrona* | N | N | N | N | N | N | Y | Y | N | ? | Y | Y | N | | Bromelicolous | Direct development |
| *Tachiramantis* sp. | N | Y | Y | Y | Y | Y | ? | Y | N | N | N | N | Y | | Lower vegetation | Direct development |
| *Serranobatrachus carmelitae* | N | N | N | N | N | N | N | N | N | N | Y | Y | N | | Riparian (boulders) | Direct development |
| *Serranobatrachus cristinae* | N | N | N | N | N | N | N | N | N | ? | Y | ? | N | | Arboreal | Direct development |
| *Serranobatrachus insignitus* | N | N | N | N | N | N | Y | Y | Y | Y | Y | Y | N | | Terrestrial | Direct development |
| *Serranobatrachus megalops/delicatus* | N | ? | ? | ? | ? | ? | Y | Y | Y | Y | Y | Y | ? | | Leaf litter, Lower vegetation | Direct development |
| *Serranobatrachus sanctaemartae* | N | Y | Y | Y | Y | Y | Y | Y | Y | Y | Y | Y | Y | | Arboreal, Lower vegetation | Direct development |
| *Geobatrachus walkeri* | N | N | N | N | N | N | N | N | N | N | Y | Y | N | | Fossorial | Direct development |
| *Rhinella horribilis* | Y | N | N | N | N | N | N | N | N | N | N | N | N | Lowland species | Terrestrial | Aquatic larvae (lotic + lentic) |
| *Boana platanera* | Y | N | N | N | N | N | N | N | N | N | N | N | N | | Arboreal | Aquatic larvae (lentic) |
| *Lithobates vaillanti* | Y | N | N | N | N | N | N | N | N | N | N | N | N | | Terrestrial, Semiaquatic | Aquatic larvae (lentic) |
| **Elevation** | 967 | 1486 | 1505 | 1524 | 1542 | 1550 | 1557 | 1577 | 1667 | 1766 | 2111 | 2459 | NA | | | |

**Protocol S1.** Considerations for RPA primer design for metabarcoding.

Given its relatively recent development, existing tools for RPA primer design, such as PrimedRPA (Higgins *et al.* 2019), are largely optimized for finding primers for single target species amplification. Metabarcoding primer design is more challenging, as the primer binding site needs to be conserved over a large range of taxa, while mismatches to non-target taxa need to be maximised. In contrast to PCR, RPA can still yield amplification with several mismatches, challenging the design of specific metabarcoding primers. While for RPA, 30-35 bp primers have been recommended, this might often require the incorporation of several degenerate bases. The latter, however, can easily lead to artefact formation and off-target amplification. Until more robust and automated tools become available, we therefore recommend manual alignment-guided design of several primer combinations and subsequent experimental validation

A simple approach to RPA metabarcoding primer design is mapping existing PCR metabarcoding primers on a target species alignment and manually extending the primer length towards one or both ends trying to minimize number of degenerate bases. Particularly at the distal positions of the primer, degenerate bases should be avoided to preclude the formation of longer fragments which cannot be easily removed with a magnetic bead clean-up. We observed that for metabarcoding, primers as short as ~22 bp in length (e.g. 16Sbr) can show good amplification in RPA. To assess amplification success during primer design, RPA products are best cleaned with magnetic beads prior to visualization as the RPA's crowding agent can prevent proper flow during gel electrophoresis. We recommend diluting RPA products 1:1 with water prior to magnetic bead clean-up as, similarly, the high density of RPA's crowding agent can prevent beads from rapidly collecting at the magnet.

**Protocol S2.** Customized protocol for field-deployable library preparation for Flongle sequencing using the SQK-LSK114 kit. RPA-based library preparation follows the supplementary protocol of Plewnia et al. (2025). Indexed RPA-products are pooled to equal amounts. Abbreviations as of SQK-LSK114 kit instructions. Mix all reagents by pipetting up and down.

1. Clean pool with magnetic beads following your manufacturer's instructions for amplicons. Generally, we recommend diluting the pool 1:1 with RNAse-free water before adding beads to prevent the RPA crowding agent from disturbing bead movement to the tube wall.

2. Measure DNA concentration using a Qubit fluorometer following the manufacturer's instructions.

3. Prepare 150-200 ng DNA for library preparation. Normalize to 24.5 µl with RNAse-free water in a PCR strip.

4. Add 0.5 µl DCS, 3.5 µl Ultra II End-prep reaction buffer and 1.5 µl Ultra II End-prep enzyme mix to your input DNA, then mix by pipetting.

5. Incubate for 30 min at ~20°C. This can be done at room temperature or (in cold environments) using a portable heating device (e.g. a 3D-printed heat bed; Plewnia et al. 2025b).

6. Incubate for 30 min at 65°C using a field-deployable heat bed.

7. Add 30 µl of resuspended AMPure XP beads and mix thoroughly by pipetting.

8. Incubate at ambient temperature for 5 min, then place on magnetic rack.

9. When all beads have pelleted on magnet, remove supernatant.

10. Wash by adding and removing 200 µl of 80% ethanol.

11. repeat 10.

12. Remove from magnet and let dry until all droplets have disappeared but avoid overdrying (fissures in bead pellet).

13. Resuspend pellet in 31 µl RNAse-free water, mix thoroughly by pipetting and incubate for 5 min.

14. Place in magnet and wait until beads have pelleted.

15. Transfer 30 µl into next well in the PCR strip.

16. Add 2.5 µl LA, 12.5 µl LNB and 5 µl Quick T4 DNA Ligase, then mix by pipetting.

17. Incubate for 10 min at room temperature (if ambient temperature is below 18°C use field-deployable heat-bed).

18. Add 20 µl resuspended AMPure XP beads, mix thoroughly by pipetting.

19. Incubate at ambient temperature for 5 min, then place on magnetic rack.

20. When all beads have pelleted on magnet, remove supernatant.

21. Wash by adding and removing 125 SFB. Make sure all beads have re-pelleted before removal of SFB as they dissolve more easily in the viscous SFB than in ethanol.

22. repeat 21.

23. Remove from magnet and let dry until all droplets have disappeared but avoid over-drying (fissures in bead pellet).

24. Resuspend pellet in 11 µl EB (this will allow two sequencing runs if needed at the cost of a single lib. prep.), mix thoroughly by pipetting and incubate for 10 min at increased temperature (e.g. in hand or heat-bed at 37°C).

25. Place in magnet and wait until beads have pelleted.

26. Transfer 5 µl each to two tubes of your PCR strip. One is intended as a backup, the other for sequencing.

27. Add 15 µl SB and 10 µl LIB to one tube containing 5µl library. Make sure to mix by pipetting right before loading.

28. In a new tube of your PCR strip, mix 117 µl of FCF with 3 µl of FCT by pipetting.

29. Check and prepare flowcell, then load it and start the sequencing run as described by the manufacturer. Load flowcell by turning the pipette plunger to avoid introduction of air. Make sure to thoroughly close all ports of the flowcell to avoid drying during the run.

**Protocol S3.** Bash code for processing of ONT sequencing data. If all programs are readily installed, this code can be run without internet connection under field conditions. The code can be run on a portable local machine. However, availability of GPU resources for basecalling strongly enhances speed of the bioinformatic workflow. Any characters in *italics* correspond to variable file names or custom thresholds that need to be replaced with the desired names/parameters.

### Basecalling with Dorado v0.9.0 in windows power shell using GPU mode (for CPU mode add `--device cpu`. Alternatively, basecalling can be done live using Dorado in MinKnow.

```
“\path\dorado\bin\dorado.exe” basecaller sup -r “\path\inputfolder” -o
“path\outputfolder” --emit-fastq
```

### Now, switch to Linux environment
### Deposit `minibar.py` script in your folder, make it executable and demultiplex with the following commands (make sure to deposit txt with index and primer data in the folder too). -T will trim off primer sequences, -F creates individual output folders. For details, check:
https://github.com/calacademy-research/minibar/tree/master

```
chmod 775 minibar.py
```

```
./minibar.py dmxLab.txt all_nanopore_files.fastq -F -T
```

### We are creating a shell-script to filter all output files at once using Seqkit. -m specifies the min length, -M the max. length of the sequence to be written, -Q is the minimum quality score. Make sure to run separate filters on environmental (short) markers and specimen barcoding (long) markers.

```
ls *.fastq | awk '{print "seqkit seq -m 160 -M 180 -Q 12 "$0" >
"gensub(/\.fastq$/, "_filtered.fastq", "g", $0)}' > filter_fastq.sh
```

```
chmod +x filter_fastq.sh
```

```
./filter_fastq.sh
```

#### **At this stage, we have to proceed differently for barcoding vs. metabarcoding data**. We first have to process specimen barcoding data and supplement our BLAST database. We therefore create consensus sequences for specimen barcodes using NGSpeciesID. Make sure to manually sort the desired files in a new folder and change your directory accordingly. The following shell script will create individual output folders.

```
ls *_filtered.fastq | awk '{split($0, a, "_filtered.fastq"); print "mkdir -
p ./NGSpeciesID/" a[1] " && NGSpeciesID --ont --consensus --medaka --fastq
\"" $0 "\" --outfolder ./NGSpeciesID/" a[1]}' > run_NGSpeciesID.sh
```

```
chmod +x run_NGSpeciesID.sh
```

```
./run_NGSpeciesID.sh
```

### Then, automatically extract your consensus sequences and name them after the sample folder using the `extract_fastas.sh` script.

### Subsequently, merge them for a quick BLAST-check for taxonomic integrity using the following command:

```
for file in *.fasta; do [[ "$file" == "name.fasta" ]] && continue; awk -v
name="${file%.fasta}" '/^>/ {print ">" name; next} {print}' "$file"; done >
name.fasta
```

### Although not mandatory, we recommend aligning and visually inspecting your consensus sequences at this stage to trim off erroneous sequence ends (NGSpeciesID is prone to create erroneous sequence overhangs beyond the actual length of your target marker). For alignment and curation, we recommend using mafft and CIAlign combined with a graphic program for visualization such as AliView or MEGA.

```
mafft –-adjustdirection name.fasta > name_aligned.fasta
```

```
CIAlign --infile name_aligned.fasta --outfile name_aligned_curated.fasta --
remove_insertions --remove_divergent
```

### Now, simply paste your sequences in fasta format into your local GenBank (In our case: the MIDORI2 subset) data (if not downloaded in fasta format you can use the `blastdbcmd` command to transform it into fasta: `blastdbcmd -db 16S -entry all -outfmt "%f" -out 16S.fasta`). In practice, this means opening the fasta e.g. in a GUI editor and pasting the new sequences at the end or using the `cat` command to concatenate two fasta files. Then make sure to manually create the header of your sequence in a similar format so that BLASTn will later provide a meaningful taxonomic annotation (i.e. headers should look like: *NoID;Eukaryota_2759;Chordata_7711;Amphibia_8292;Anura_8342;Bufonidae_8382;Atelopus_47578;Atelopus_laetissimus_consensus*). Then, create a BLAST database from your supplemented fasta file:

```
makeblastdb -in name.fasta -dbtype nucl -out dbname
```

### Check if BLAST can access the database.

```
blastdbcmd -db dbname -info
```

### Now you can get back to your metabarcoding data, starting from the filtered fastq files. To reduce file size, change .fastq to .fas

```
ls *filtered.fastq | awk '{print "fastq_to_fasta -i \"" $0 "\" -o \""
substr($0, 1, length($0)-6) ".fas\" -n"}' > fas.sh
```

```
chmod +x fas.sh
```

```
./fas.sh
```

### Now, dereplicate sequences

```
ls *filtered.fas | awk '{print "vsearch --derep_fulllength \"" $0 "\" --
output \"" substr($0, 1, length($0)-4) ".derep\" --sizeout --relabel Uniq"}'
> derep.sh
```

```
chmod +x derep.sh
```

```
./derep.sh
```

### Instead of further clustering or denoising approaches, we recommend the direct-BLAST approach (Plewnia et al. 2025) for ONT data given that sequence error distance can exceed genetic distance between closely related species.

```
ls *.derep | awk '{print "blastn -db /path/dbname -query \"" $0 "\" -
max_target_seqs 1 -out \"" substr($0, 1, length($0)-4) "_OTUs.out\" -outfmt
```

```
\"6 qseqid sseqid stitle pident length mismatch gapopen qstart qend sstart send evalue bitscore staxids\""}' > blast.sh

chmod +x blast.sh

./blast.sh
```

### The output can be read and processed with any GUI text editor and contains row-wise taxonomic annotations for each unique sequence. To generate an OTU table, weighted (by size of dereplication) sums can be built per taxon name using custom R code. An example can be found below in protocol S4.

**Protocol S4**. Example R script for downstream analysis of eDNA metabarcoding results, summarizing OTU/species read counts for all samples (based on the OTU tables with BLAST results as generated and formatted in protocol S3). Characters in *italics* correspond to column names or custom thresholds that need to be replaced with the desired names/parameters. Note that, depending on the formatting of the taxonomy string, redundant symbols may persist in the taxon names, and a critical evaluation of the final results is required.

```
# Save and place script (.R file) in folder with all input OTU tables; set working directory to script location:
setwd(dirname(rstudioapi::getActiveDocumentContext()$path))

# Load Tidyverse (version 2.0.0 used for the script below):
library(tidyverse)

# List all OTU tables (named as .out files; see protocol S3) in the folder:
file_list <- list.files(pattern = "\\.out$")

### Process all OTU tables and return summarized read counts for each OTU/species, also retaining family
designation. Numbers (#1, #2, #3) require specific column input (see more information below the script).
## Read the files (Here: as tab-delimited tables)
## Extracting & cleaning taxonomic data:
# — in this example: keep only records with a sequence similarity ≥ 95%
# — for sequence header containing "size=N"; extract N as numeric (read) count
# — extract family name from the taxonomy string (filtering for "dae" word ending)
# — extract species name (assuming they are the last semicolon-delimited field)
# — remove ‘noise’ extracted with the species and family names from the taxonomy string
# — replace the first underscore with a space ("Genus_species" → "Genus species")
# — remove remaining underscores, drop remaining digits
## Sum all read counts per OTU/species, with each column summarizing one input OTU table.

process_file <- function(file_name) {
 data <- read_tsv(file_name, col_names = FALSE)

 cleaned <- data %>%
  filter(column >= 95) %>% # (1)
  mutate(column = as.numeric(str_extract(column, "(?<=size=)\\d+")), # (2)
   family = str_extract(column, "\\w*dae"), # (3)
   family = str_remove(family, "^family_"),
   family = str_remove(family, "_\\d+$"),
   species = str_extract(column, "[^;]+$"), # (3)
   species = species %>%
    str_remove_all("(?i)^species_") %>%
    str_remove_all("_consensus") %>%
    str_remove_all("(?i)[sS][pP][0-9]*") %>%
    str_replace("_", " ") %>%
    str_remove_all("_") %>%
    str_remove_all("\\d+")) %>%

  select(family, species, count = column) # (2)

 summarized <- cleaned %>%
  group_by(family, species) %>%
  summarise(!!file_name := sum(count, na.rm = TRUE), .groups = "drop")

 return(summarized)
}
```

### (1) Provide column containing sequence identity (% sequence similarity).
### (2) Provide column containing the total read counts per OTU (specified as "size=N" after unique identification number as in our example).
### (3) Provide column with full taxonomy string (formatted e.g. *NoID;Eukaryota_2759;Chordata_7711;Amphibia_8292;Anura_8342;Bufonidae_8382;Atelopus_47578;Atelopus_laetissimus_consensus*) as in our example.

##### Collate and clean all data:
### — Replace NAs (species absent in some files) with 0 across columns
### — Set all values ≤ 5 to 0 (regarding read counts below this threshold as sequencing noise)
### — Remove taxa rows where the sum across all samples is 0 (after filtering)

```
all_data <- map(file_list, process_file) %>%
  reduce(full_join, by = c("family", "species")) %>%
  mutate(across(-c(family, species), ~replace_na(.x, 0))) %>%
  mutate(across(-c(family, species), ~ifelse(.x <= 5, 0, .x))) %>%
  filter(rowSums(across(-c(family, species))) > 0)
```

### Save final summary table

```
write_tsv(all_data, "summary_table.txt")
```
